# An evolutionary resolution of manipulation conflict

**DOI:** 10.1101/003707

**Authors:** Mauricio González-Forero

## Abstract

Individuals can manipulate the behavior of social partners. However, manipulation may conflict with the fitness interests of the manipulated individuals. Manipulated individuals can then be favored to resist manipulation, possibly reducing or eliminating the manipulated behavior in the long run. I use a mathematical model to show that conflicts where manipulation and resistance coevolve can disappear as a result of the coevolutionary process. I find that while manipulated individuals are selected to resist, they can simultaneously be favored to express the manipulated behavior at higher efficiency (i.e., providing increasing fitness effects to recipients of the manipulated behavior). Efficiency can increase to a point at which selection for resistance disappears. This process yields an efficient social behavior that is induced by social partners, and over which the inducing and induced individuals are no longer in conflict. A necessary factor is costly inefficiency. I develop the model to address the evolution of advanced eusociality via maternal manipulation (AEMM). The model predicts AEMM to be particularly likely in taxa with ancestrally imperfect resistance to maternal manipulation. Costly inefficiency occurs if the cost of delayed dispersal is larger than the benefit of exploiting the maternal patch. I discuss broader implications of the process.

## Introduction

In some taxa, individuals can control partially or completely the behavior of other individuals, an action referred to as manipulation (Alexander, 1974, Dawkins, 1978). For example, baculoviruses manipulate their host, a moth caterpillar, to climb trees; the caterpillars then die and liquefy at the tree top causing a “virus rain” in the foliage below, thereby facilitating infection of new hosts (Hoover *et al.*, 2011). Workers in social insects can induce their siblings to develop as workers or queens by adjusting their siblings’ nutrition (Wheeler, 1986, O’Donnell, 1998). *Drosophila* males manipulate their sexual partners by transferring seminal proteins during mating (Wolfner, 2002). Manipulation is facilitated when an individual has direct access to another individual’s physiology, as is the case for internal parasites (Hughes *et al.*, 2012, Adamo and Webster, 2013), for parents and offspring (Haig, 1993), and for mating partners (Arnqvist and Rowe, 2005). In the absence of direct access to another individual’s physiology, an individual may manipulate another one through coercion, sensory exploitation, deception, and self-deception. In particular, dominant individuals may coerce subordinates into helping roles (Clutton-Brock and Parker, 1995), males may stimulate females’ pre-existing preferences to induce mating (Holland and Rice, 1998), and humans may deceive themselves to fool social partners into behaving in a given fashion (Trivers, 2011).

Manipulation can give rise to unlikely behaviors because the costs of expressing the behaviors are not paid by the manipulators, but by the subjects of manipulation (or “subjects” for short). As a result, costly behaviors can evolve under less stringent conditions (i.e., smaller benefit-cost ratios) than if the behaviors were performed spontaneously; that is, without manipulation (Alexander, 1974, Trivers, 1974, Charlesworth, 1978). However, costly behaviors diminish the reproductive success of the subjects. Resistance to manipulation is then favored if resistance is less expensive than accepting manipulation (Pagel *et al.*, 1998). Manipulators and subjects can thus disagree in their preferred expression level of the manipulated behavior, which constitutes an evolutionary conflict (Trivers, 1974).

Evolutionary conflicts can have diverse results. Mathematical theory indicates that a manipulation conflict can yield at least four possible outcomes. First, the complete victory of resistance where the manipulated behavior is eliminated. Second, the complete victory of manipulation where the manipulated behavior is fully maintained. Third, an intermediate behavior between the favored outcomes of the two parties. And fourth, perpetual cycles between high and low manipulation and resistance (e.g., Parker and Macnair, 1979, Robert *et al.*, 1999, Gavrilets *et al.*, 2001). Which outcome is reached depends on the magnitude and nature of the costs paid by each party (Godfray, 1995, Clutton-Brock, 1998, Uller and Pen, 2011), the initial conditions, and the relative genetic variances of manipulation and resistance (Gavrilets, 2000, Gavrilets and Hayashi, 2006).

As the evolution of manipulation and resistance proceeds, the nature of the conflict can change. In particular, the costs and benefits of the manipulated behavior can evolve if they have a genetic basis (Charlesworth, 1978, Worden and Levin, 2007, Akçay and Roughgarden, 2011). A genetic basis for the costs and benefits of a manipulated behavior is possible because they depend on the extent with which the manipulated behavior is expressed, which can be controlled by manipulators and subjects of manipulation. The evolution of costs and benefits could then increase or decrease the level of conflict. As a result, the outcome of a conflict can be substantially different from what it would be if costs and benefits are taken as constants. Here I ask what the evolution of fitness payoffs can do to the outcome of a manipulation conflict.

I show that the manipulation conflict can disappear as a result of the evolution of payoffs released by manipulation. This conflict resolution brings the interests of the subjects of manipulation to match those of the manipulator. The reason is that manipulation not only favors the evolution of resistance, but also the evolution of the efficiency with which the manipulated behavior is performed. If the efficiency of the manipulated behavior becomes sufficiently high, resistance to manipulation becomes disfavored. Because the conflict is eliminated, I refer to the resulting behavior as being induced rather than manipulated. The result is an efficient induced behavior over which inducing and induced partners do not conflict. To show this, I develop a mathematical model of maternal manipulation where offspring are manipulated to stay in the maternal patch. As offspring evolve resistance to manipulation, they also become efficient helpers giving large fitness benefits to siblings. The outcome is offspring that are 1) maternally induced to stay in the maternal patch, 2) that are efficient helpers, and 3) that are not in conflict with their mother over their helping role. These three items match defining features of advanced eusociality, where workers are maternally induced into worker roles, can be highly specialized to perform tasks, and show relatively little conflict over their helping role (Wilson, 1971, Michener, 1974, Sherman *et al.*, 1991, Crespi and Yanega, 1995, Hölldobler and Wilson, 2009, Bignell *et al.*, 2011). The model predicts that advanced eusociality arising from this process requires ancestrally imperfect resistance probability and ancestral inefficiency costs.

## Model

Consider a finite population of sexual individuals with deterministic reproduction, so genetic drift is ignored. The genetic system can be diploid or haplodiploid. The population is distributed in an area of a fixed size which is subdivided into patches, all of approximately the same size. In each patch, one singly mated female and possibly her mating partner gather resources for reproduction. The amount of resources they gather is proportional to the patch size. The mated female produces offspring, the number of which is proportional to the amount of resources gathered. So offspring number is proportional to the patch size. The average patch size decreases as the population increases, and increases as the population decreases. Hence, the population size remains constant.

The mother produces offspring in two subsequent broods. The first brood reaches adulthood while the second brood is not yet mature. The mother and possibly the father provide parental care to both broods. Once the second brood reaches adulthood, the parents die. After each brood reaches adulthood, the brood disperses from the maternal patch to a common mating pool. All individuals in the mating pool mate once and randomly. Then, each mated female colonizes a random patch, possibly together with her mating partner, and the cycle starts anew. Competition for patch size is thus global.

Maternal manipulation and offspring resistance are allowed to occur. A focal mother manipulates offspring by attempting to delay the dispersal of the first brood with probability *p*_*m*_, so that first-brood offspring stay in the maternal patch for a fraction of their adulthood. I make the simplifying assumption that the mother manipulates both sexes equally. A manipulated first-brood individual resists with probability *q*_1_ and leaves the maternal patch without delay. Alternatively, a manipulated first-brood individual acquiesces (i.e., does not resist) with probability 1 − *q*_1_ and stays in the maternal patch for some portion of its adulthood. An acquiescing (i.e., delayed) individual expresses parental care with probability *y*_1_ while in the maternal patch. I also make the simplifying assumption that the probability of expressing parental care is equal for acquiescing individuals of either sex. This alloparental care is directed randomly to the available brood (i.e., the second one). I refer to *y*_1_ as helping efficiency. I assume manipulation *p*_*m*_, resistance *q*_1_, and helping efficiency *y*_1_ to be uncorrelated quantitative genetic traits. The population average values of manipulation, resistance, and helping efficiency are *p*, *q*, and *y* respectively. The three decisions individuals can make are illustrated in Fig. 1.

**Figure 1:**
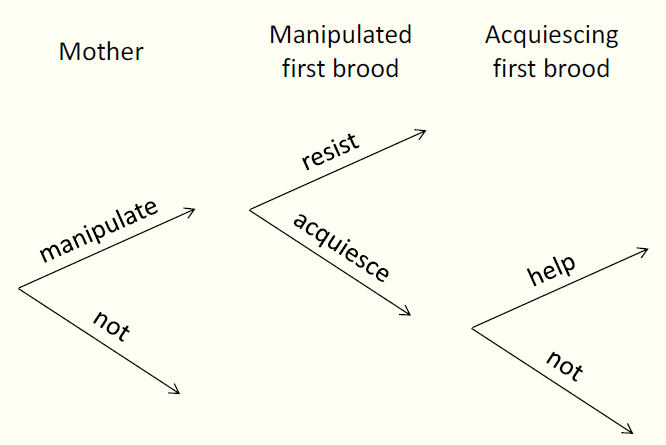
The three decisions individuals can make. Mothers manipulate with probability *p* first-brood offspring to stay as adults. Manipulated first-brood offspring resist with probability *q* and leave without delay. Otherwise, they acquiesce with probability 1 − *q* and stay for some period. Acquiescing individuals help with probability *y* to raise the second brood.

Manipulation is assumed to be executed in a way that does not affect the condition of the subjects of manipulation, and that does not affect the ability of the mother to produce the second brood. Thus, I assume both resistance and manipulation to be costless. These assumptions may hold for instance if manipulation is done via cheap pheromones that first-brood individuals can block with little direct fitness costs. The assumptions of costless manipulation and resistance lead to a simpler model, and highlight that the evolution of acquiescence does not require resistance costs (for an evaluation of the effect of resistance and manipulation costs on the evolution of manipulated behaviors, see González-Forero and Gavrilets (2013)). However, manipulation affects the ability of acquiescing individuals to become parents themselves. First, regardless of whether an acquiescing individual helps, this individual has a reduced probability of becoming a parent if delayed dispersal translates into missed reproductive opportunities or less time to start a new nest. Second, if an acquiescing individual helps, a reduced probability of becoming a parent arises if by helping, the individual spends energy necessary for its own dispersal and reproduction. In contrast, if an acquiescing individual does not help, it can exploit the resources of the maternal patch for its own benefit, thereby increasing its potential to become a parent itself.

These fitness payoffs are modeled as follows. The reduction in the probability that a delayed individual becomes a parent, independently of whether the delayed individual helps, is denoted by *c*_*d*_. The additional reduction in the probability that a delayed individual becomes a parent due to helping is *c*_*h*_. On the other hand, the increase in the probability of becoming a parent due to the exploitation of the maternal patch while not helping is *b*_*e*_. For simplicity, I ignore any frequency dependence in the payoffs *c*_*d*_, *c*_*h*_, and *b*_*e*_, and I treat them as constant. The total cost of acquiescence for a focal delayed first-brood individual is thus equal to

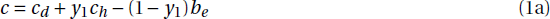

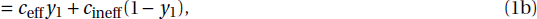

where the cost of efficiency and inefficiency are defined as

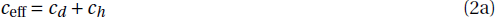

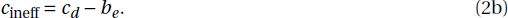

In this paper, I report the behavior of the model when there is a cost of inefficiency (*c*_ineff_ > 0), so I assume throughout that the cost of delayed dispersal is greater than the benefit of exploiting the maternal patch (*c*_*d*_ > *b*_*e*_).

It remains to account for the fitness effects of manipulation on the second brood. A delayed first-brood individual that helps increases the survival of recipient second-brood offspring. The increase in survival received by a random second-brood recipient is

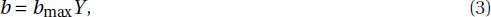

where *Y* is the average helping probability among delayed individuals in the patch, and *b*_max_ is the benefit a recipient of help gets when all delayed individuals in the patch help at their maximum efficiency. Denoting by *s*_0_ the baseline probability of becoming a parent (i.e., the probability that offspring become parents when manipulation does not occur), I let *b*_max_ = 1 − *s*_0_.

I follow the methods of Taylor and Frank (1996) and Frank (1997) to obtain dynamic equations for the coevolution of manipulation *p*, resistance *q*, and helping efficiency *y* (see Appendix). At any given time the population is divided into three classes of individuals: mothers, first-brood individuals, and second-brood individuals. This treatment yields three regression relatednesses that affect the evolutionary dynamics: the relatedness *ρ*_21_ of first-brood offspring toward second-brood offspring, and those of the mother toward the first and second brood (*ρ*_1*m*_ and *ρ*_2*m*_ respectively). For class-structured populations, the direction of evolutionary change usually depends on regression relatedness weighted by the individual reproductive value of the recipient over that of actor, which is called life-for-life relatedness (Hamilton, 1972, Bulmer, 1994). However, here the direction of evolutionary change is found to be determined by regression relatednesses weighted by equilibrium class frequencies rather than by individual reproductive values. The weighting by class equilibrium frequencies arises because the evolving traits affect survival rather than fertility. Thus, the dynamics are in terms of the equilibrium relatednesses *r*_*ji*_ of actor *i* toward recipient *j*, which are defined as *r*_*ji*_ = *ρ*_*ji*_*u*_*j*_/*u*_*i*_. The quantities *u*_*i*_ and *u*_*j*_ are the equilibrium frequency of individuals of class *i* and *j* respectively. For simplicity, I drop the subscripts for the relatedness of first-brood offspring toward second-brood offspring and write *ρ* = *ρ*_21_, and *r* = *r*_21_.

## Results

### Numerical illustration

The coevolution of manipulation, resistance, and helping efficiency can change the direction of selection for resistance. For illustration, suppose that the population is diploid. Thus, with the assumptions stated in the model section, the regression relatednesses of mother-to-offspring and of first-to second-brood offspring are *ρ*_1*m*_ = *ρ*_2*m*_ = *ρ* = 1/2 (Bulmer, 1994, Roze and Rousset, 2004). Numerical solutions for manipulation *p*, resistance *q*, and helping efficiency *y* from the dynamic equations (19) are shown in Fig. 2. Both maternal manipulation and offspring resistance are favored at the start of the process in Fig. 2. In Fig. 2A there is no genetic variation for helping efficiency, which then cannot evolve. In this case, resistance eliminates the manipulated behavior and all first-brood offspring disperse upon reaching adulthood (i.e., the attained equilibrium is *p*^*^ (1 − *q*^*^) = 0). In Fig. 2B genetic variation for helping efficiency is present. In this case, helping efficiency increases over time although individuals are initially disfavored to stay in the maternal patch. After around thirty thousand generations, helping efficiency becomes high enough that first-brood individuals become favored to stay in the maternal patch. The outcome is that mothers cause all first-brood individuals to stay, first-brood individuals acquiesce, and help at their maximum efficiency.

**Figure 2:**
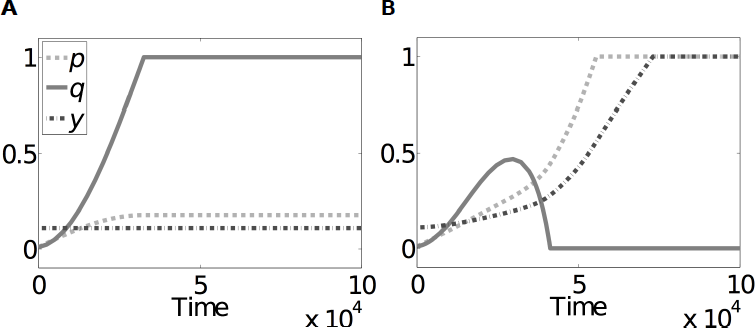
Coevolution of manipulation *p*, resistance *q*, and helping efficiency *y*. Numerical solutions of eqs. (19) are shown. (A) There is no genetic variation for helping efficiency (*V*_*y*_ = 0). Resistance evolves and eliminates the manipulated behavior (i.e., the probability that first-brood offspring stay in the maternal patch is *p*^*^ (1 − *q*^*^) = 0 at the end). (B) Same conditions as in (A), but there is genetic variation for helping efficiency (*V*_*y*_ = 0.001). Helping efficiency increases and after ≈ 30 × 10^3^ generations, resistance decreases and is eliminated. The remaining parameter values for both panels are *p*_0_ = *q*_0_ = 0.01, *y*_0_ = 0.11, *ρ* = *ρ*_1*m*_ = *ρ*_2*m*_ = 1/2, *V*_*p*_ = 0.001, *V*_*q*_ = 0.1, *α* = *σ* = *η* = *s*_0_ = 1/2, *c*_eff_ = 0.2, and *c*_ineff_ = 0.012.

This coevolutionary process eliminates the mother-offspring conflict over offspring dispersal. Throughout the process, the mother’s inclusive fitness through maternal manipulation is maximized at zero offspring resistance (Fig. 3A). In contrast, first-brood offspring’s inclusive fitness through resistance is initially maximized at full resistance, but the slope of their inclusive fitness gradually changes from positive to negative (Fig. 3B). The change in slope of offspring’s inclusive fitness through resistance renders this inclusive fitness maximized at zero resistance, thereby eliminating the mother-offspring conflict (Fig. 3C). Because of the lack of conflict, I refer to the final maternally triggered behavior as being induced rather than being manipulated.

**Figure 3:**
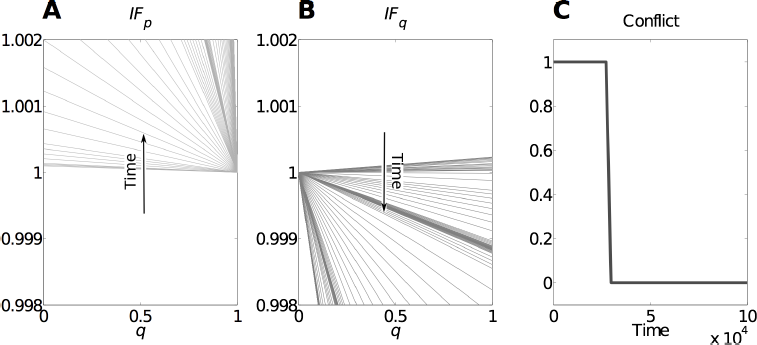
Evolutionary resolution of manipulation conflict. (A) Mother’s inclusive fitness through manipulation *IF*_*p*_ vs. possible values of resistance *q* during the process in Fig. 2B. The lowest line is mother’s inclusive fitness at time 1 in Fig. 2B, and the lines further up correspond to mother’s inclusive fitness as time increases. (B) First-brood offspring’s inclusive fitness through resistance *IF*_*q*_ vs. possible values of resistance during the process in Fig. 2B. The highest line is offspring’s inclusive fitness at time 1 in Fig. 2B, and the lines further down correspond to offspring’s inclusive fitness as time increases. In (A), the optimum inclusive fitness for the mother is at *q* = 0 throughout, while in (B) the optimum inclusive fitness for first-brood offspring is initially at *q* = 1 and later at *q* = 0. (C) The level of conflict over time. The level of conflict is the distance between the preferred trait values of the two parties. After ≈ 30 × 10^3^ generations, the conflict disappears. For the three panels, the same parameter values are used as in Fig. 2B. The inclusive fitness through trait *i* (= *p*, *q*) is *IF*_*i*_ = *IF*_0_ + *ih*_*i*_, where the baseline inclusive fitness (*IF*_0_) is set to 1, and the inclusive fitness effect of trait *i* (*h*_*i*_) is given by the right-hand side of (eqs. (19a) or ((19b) divided by *V*_*i*_ respectively. The level of conflict is *C* = |max_*q*_(*IF*_*p*_) − max_*q*_(*IF*_*q*_)|, where max_*q*_(*x*) gives the resistance *q* that maximizes *x*.

### Evolutionary change in each trait

The population-average manipulation *p*, resistance *q*, and helping efficiency *y* increase respectively (see eqs. (19) in the Appendix) when

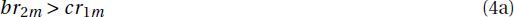

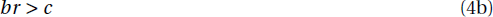

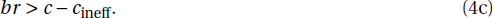

Manipulation, acquiescence, and helping efficiency are each favored when their respective inclusive fitness effect is positive (conditions (4)). Manipulation conflict occurs when manipulation is favored but acquiescence is not (i.e., when in eq. (4a) is met but in eq. (4b) is not). In that case, mothers attempt to delay first-brood offspring in the maternal patch against the latter’s inclusive fitness interests. Offspring can rebel against manipulation by either resisting (i.e., dispersing from the maternal patch) or by refusing to help. The conditions for the evolution of these two forms of rebellion are different if the cost of inefficiency *c*_ineff_ is not zero (see in eqs. (4b) and (4c)). The different conditions for the evolution of acquiescence and helping efficiency can cause conflicting selection within first-brood offspring. Thus, helping efficiency may evolve even though acquiescence is not favored.

Conditions (4) do not specify the conditions for conflict resolution because the benefit *b* and cost *c* evolve as helping efficiency *y* changes. Consequently, whether or not conditions (4) are met varies with the evolution of helping efficiency. In order to determine the conditions for conflict resolution, a dynamic analysis is necessary (see §2 in the online Supporting Information (SI)).

### Conditions for conflict resolution

The system evolves either to a state where manipulation disappears (*p*^*^ = 0), to a state where resistance is complete (*q*^*^ = 1), or to induced behavior where manipulation, acquiescence, and helping efficiency are established [(*p*^*^, *q*^*^, *y*^*^) = (1, 0, 1)].

The evolution of induced behavior requires two conditions regarding resistance. First, acquiescence must be favored when first-brood offspring help at their maximum efficiency, which occurs if

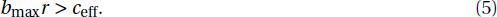

When condition (5) holds, the coevolutionary dynamics of resistance *q* and helping efficiency *y* are as described in Fig. 4A. Acquiescence can be disfavored at the start of the process, and the evolution of helping efficiency can render acquiescence favored if the population starts in the dark gray area in Fig. 4A. The population starts in the either the gray or dark gray area in Fig. 4A if the next condition is met. Second, induced behavior requires that the probability of resistance is initially small enough, which occurs if

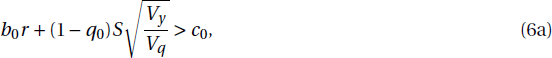

where

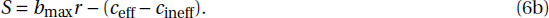

The variables with subscript “0” refer to the value of the variable at the initial time. The quantity *S* measures selection for helping efficiency, which is positive when condition (5) holds. *V*_*q*_ and *V*_*y*_ are the additive genetic variances for resistance and helping efficiency respectively.

**Figure 4:**
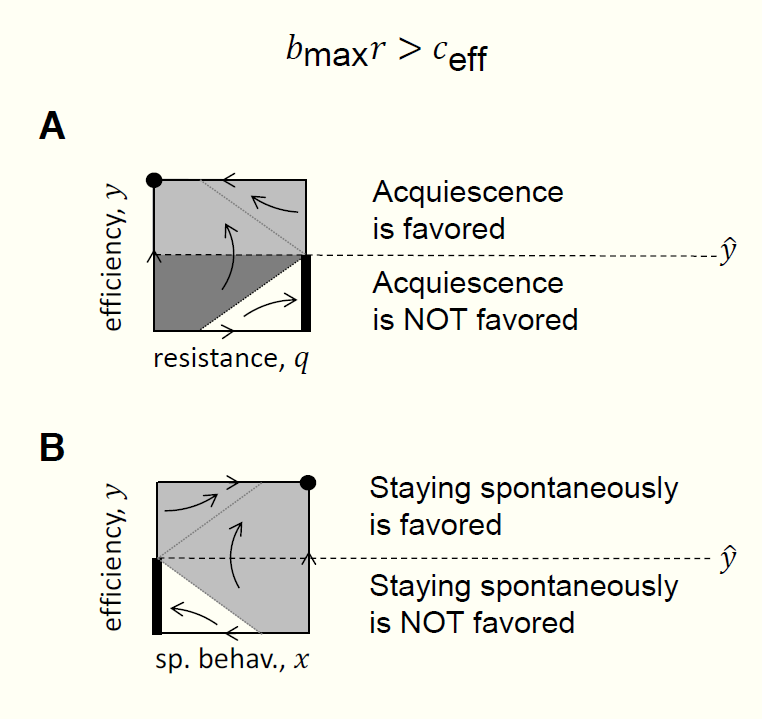
Coevolutionary dynamics of resistance *q* and spontaneous behavior *x* with helping efficiency *y* when *b*_max_*r* > *c*_eff_. The arrows indicate the direction of change (the arrows at the boundaries indicate the partial change with respect to the direction of the boundary). Thick strokes indicate stable equilibria. (A) Coevolution of resistance *q* and helping efficiency *y*. Acquiescence is disfavored below the dashed line and is favored above it. The dashed line is the critical helping efficiency *ŷ* = *c*_ineff_/*S* (obtained from in eq. (4b)). If the population starts in the gray areas, it converges to acquiescence and maximum helping efficiency (large dot). Thus, for final acquiescence, acquiescence need not be favored initially if the probability of resistance is initially sufficiently small (i.e., if the population starts in the dark gray area). (B) Coevolution of spontaneous behavior *x* and helping efficiency *y*. If the population starts in the gray area, it converges to spontaneous behavior and maximum helping efficiency (large dot).

Condition (6a) is related to Hamilton’s rule (Hamilton, 1964, 1970). Hamilton’s rule states that acquiescence is favored at the initial time if *b*_0_*r* > *c*_0_ (from in eq. (4b)). The additional term in condition (6a) measures the speed of increase in helping efficiency relative to that of resistance 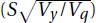 and the opportunity that helping efficiency has to render acquiescence favored (1 − *q*_0_). Because this additional term is positive when in eq. (5) holds, condition (6a) requires less stringent conditions (smaller *b*/*c* ratios) to be met than those required for acquiescence to be favored at the initial time (*b*_0_*r* > *c*_0_). Condition (6a) may then be seen as defining a relaxed Hamilton’s rule, which rather than giving the direction of selection specifies when acquiescence can be obtained in the long run.

The evolution of induced behavior also requires two conditions regarding manipulation. First, manipulation must be favored when first-brood offspring help at their maximum efficiency (in eq. (S25a) in the SI). Second, the evolution of helping efficiency must be able to render manipulation favored (in eq. (S25c) in the SI). If the probability of manipulation is initially small, the second condition regarding manipulation simply states that manipulation must be favored initially.

Four conditions are then necessary and sufficient for induced behavior (in eqs. (S25) in the SI). If manipulation *p* and resistance *q* are initially small, induced behavior (*p*^*^, *q*^*^, *y*^*^) = (1, 0, 1) evolves if all the following conditions hold:

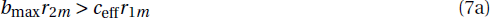

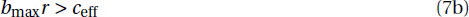

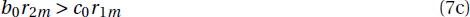

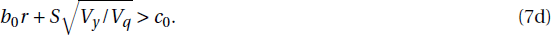

Conditions (7a) and (7b) respectively state that both manipulation and acquiescence must be favored when helping efficiency is maximal; condition (7c) states that manipulation must be initially favored; and condition (7d) guarantees that acquiescence becomes favored as the population evolves.

The evolutionary resolution of manipulation conflict occurs when induced behavior is obtained and acquiescence is not initially favored (i.e., conditions (7) are met but condition (4b) is not met initially). The region of parameter space in which the conflict is resolved is narrow (black regions in Fig. 5).

**Figure 5:**
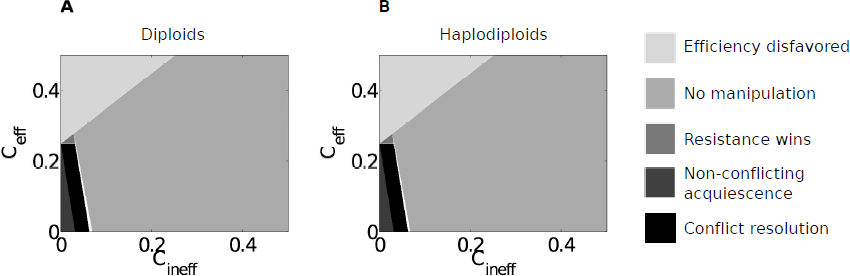
Conflict resolution across values of efficiency and inefficiency costs. (A) Diploids. (B) Haplodiploids. For both panels, in the lightest gray area, helping efficiency is disfavored. For the other shades, helping efficiency is favored. In addition, for light gray: manipulation does not evolve; for gray: manipulation evolves, but resistance wins; for dark gray: manipulation evolves but acquiescence is favored from the start and it is established at the end; and for black: manipulation and resistance evolve, but resistance is eliminated by the evolution of helping efficiency. Specifically, each region satisfies the following. For the lightest gray, *y*(0) > *y*(1); for the other shades *y*(0) < *y*(1). In addition, light gray: *p*(0) > *p*(1), and *p*(end) < 0.1; gray: *p*(0) < *p*(1), *q*(0) < *q*(1), and *q*(end) > 0.9; dark gray: *p*(0) < *p*(1), *q*(0) > *q*(1), and *q*(end) < 0.1; and black: *p*(0) < *p*(1), *q*(0) < *q*(1), *y*(0) < *y*(1), *p*(end) > 0.9, *y* (end) > 0.9, and *q*(end) < 0.1. White areas do not satisfy any of these conditions. The end is at 10^6^ generations. Parameter values are as in Fig. 2 except that *V*_*p*_, *V*_*y*_, *V*_*y*_ = 0.01 and in (B) *η*_♀_ = 1/2, *η*_♂_ = 1, *η* = *ση*_♀_ + (1 − *σ*)*η*_♂_ = 3/4, *ρ* = *σ*[*σ*3/4 + (1 − *σ*)1/2] + (1 − *σ*) [*σ*/4 + (1 − *σ*)/2] = 1/2, and *ρ*_1*m*_ = *ρ*_2*m*_ = *σ*/2 + (1 − *σ*) = 3/4 (regression relatedness values are taken from Bulmer (1994)).

However, the region for conflict resolution can be wider than the region in which first-brood offspring are favored to stay from the beginning of the process (i.e., non-conflicting acquiescence; dark gray regions in Fig. 5). In this simple model, where the mother equally manipulates both sexes, both sexes are equally efficient, and the sex ratio is equal in both broods, the region of conflict resolution can be the same for both diploids and haplodiploids (Fig. 5).

### Spontaneous behavior

The evolution of helping efficiency could render spontaneous (i.e., unmanipulated) helping favored just as it does for induced behavior. In §1 of the SI, I build an analogous model in which the probability *x*_1_that a first-brood individual stays in the maternal patch is fully under control of the staying individual. A spontaneously staying first-brood individual expresses alloparental care toward the second brood with probability *y*_1_. The population averages of the spontaneous behavior and helping efficiency are *x* and *y* respectively.

The coevolutionary dynamics of the spontaneous behavior *x* and helping efficiency *y* are a mirror image of those of resistance *q* and helping efficiency *y* (Fig. 4B). As a result, if staying spontaneously is initially disfavored, the evolution of helping efficiency can render it favored. Two conditions must be satisfied for efficient spontaneous behavior to be obtained [(*x*^*^, *y*^*^) = (1, 1)]. First, spontaneous behavior must be favored when helping efficiency is maximal (same condition (5) as for acquiescence). Second, the probability of staying spontaneously must be initially *large* enough (condition (6a) after changing 1 − *q*_0_ for *x*_0_ and *V*_*q*_ for the additive genetic variance for staying spontaneously *V*_*x*_). Hence, the opportunity for helping efficiency to render spontaneous behavior favored is *x*_0_. If the initial probability *x*_0_ of staying spontaneously is small (as is expected to be the case for altruistic traits), then the second condition for efficient spontaneous behavior simply requires that the spontaneous behavior is favored at the initial time. Hence, if the ancestral probability of spontaneously staying is small, the evolution of helping efficiency cannot not render the spontaneous behavior favored if it is not favored initially.

Consequently, induced behavior (*p*^*^, *q*^*^, *y*^*^) = (1, 0, 1) can be obtained under less stringent conditions (smaller initial *b*/*c* ratios) than spontaneous behavior (*x*^*^, *y*^*^) = (1, 1). In particular, suppose that the initial benefit *b*_0_ and cost *c*_0_ are the same under manipulated and spontaneous behavior. Assume also that the relatedness *r* of first-to second brood is the same under manipulated and spontaneous behavior. Then, if manipulation, resistance, and staying spontaneously are all initially unlikely (i.e., *p*_0_, *q*_0_, *x*_0_ ≈ 0), induced behavior can be obtained (condition (7d) is met) while spontaneous behavior is not obtained (*b*_0_*r* < *c*_0_) if

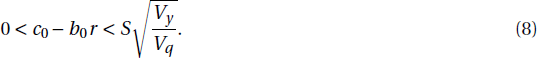

Condition (8) specifies when induced behavior can be expected but spontaneous behavior fails to evolve. This condition summarizes that the evolution of helping efficiency allows induced behavior to require less stringent conditions than spontaneous behavior since condition (8) cannot be satisfied if helping efficiency cannot evolve (i.e., if *SV*_*y*_ = 0).

Because induced and spontaneous behavior can evolve under different conditions, predictions may be derived to test whether or not advanced eusociality in a given taxon is the result of manipulation.

### Discerning whether advanced eusociality stems from manipulation

The ancestral conditions give a distinction between induced and spontaneous behavior. Induced behavior requires ancestrally imperfect resistance probability (black line in Fig. 6A). Under the same ecological conditions and if the ancestral benefit *b*_0_ and the ancestral cost *c*_0_ are the same under manipulated and spontaneous behavior, spontaneous behavior requires a sufficiently large ancestral probability of staying spontaneously (dashed gray line in Fig. 6A).

**Figure 6:**
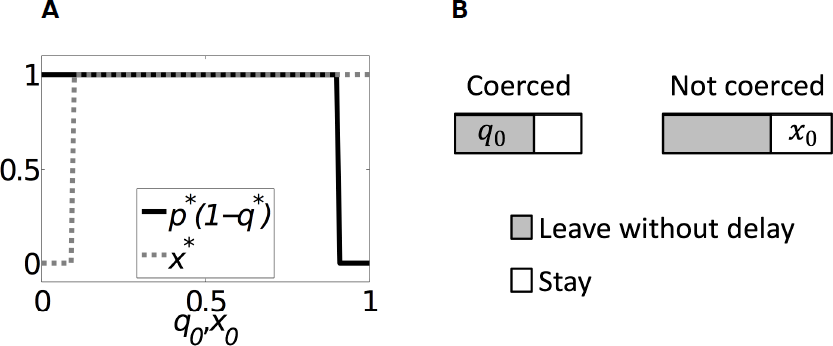
Discerning between manipulation and spontaneous helping. (A) Equilibrium values for induced and spontaneous behavior. The black line shows the predicted fraction of first brood that stay under manipulation vs. the ancestral resistance probability (*q*_0_). The dashed gray line shows the predicted fraction of first brood that stay under spontaneous behavior vs. the ancestral probability of staying spontaneously (*x*_0_). For advanced eusociality from manipulation, the ancestral resistance probability (*q*_0_) must be small enough. In contrast, for advanced eusociality from spontaneous behavior and the same ecological conditions, the ancestral probability of staying spontaneously (*x*_0_) must be large enough. Parameter values are as in Fig. 2 except that *V*_*p*_, *V*_*q*_, *V*_*x*_, *V*_*y*_ = 0.01, *c*_eff_ = 0.06, and *c*_ineff_ = 0.05. (B) Estimation of the ancestral probabilities of resistance (*q*_0_) and of staying spontaneously (*x*_0_). A fraction of the first-brood individuals in the ancestral population is maternally coerced. The ancestral probability of resistance (*q*_0_) is given by the fraction of coerced first brood that leave without delay. The ancestral probability of staying spontaneously (*x*_0_) is given by the fraction of non-coerced first brood that stay in the maternal patch for a sufficiently large portion of their adulthood so that their reproductive success is decreased.

Although it is not possible to directly determine ancestral conditions except when experimental evolution is feasible, indirect estimation of ancestral conditions may be possible. Consider an advanced eusocial population A of interest. Assume there is an extant population B satisfying the following requirements. 1) The population B is not advanced eusocial; 2) it is very close phylogenetically to population A and it has not been exposed to the manipulation mechanism that could have brought population A to advanced eusociality; and 3) it has the following life-history properties: offspring are produced in two subsequent broods, first-brood individuals are maternally manipulated in a detectable way (e.g., via coercion), and some of the first-brood offspring stay as adults in the maternal patch. Then, the ancestral probability of resistance (*q*_0_) and of staying spontaneously (*x*_0_) for population A can be estimated in population B (Fig. 6B). The ancestral probability of resistance (*q*_0_) is given by the fraction of the manipulated first brood that leave the maternal patch (gray area on the left side of Fig. 6B). In contrast, the ancestral probability of spontaneously staying (*x*_0_) corresponds to the fraction of the first brood that stay without being manipulated (white area on the right side of Fig. 6B).

A large resistance probability in population B rejects the hypothesis that the advanced eusociality in population A arose from the resolution of a conflict caused by the manipulation mechanism evaluated in B. An imperfect resistance probability in B is consistent with advanced eusociality via resolution of manipulation conflict in A (black line in Fig. 6A). Similarly, a small probability of staying spontaneously in population B rejects the hypothesis that the advanced eusociality in population A arose because the evolution of helping efficiency rendered spontaneous behavior favored. A substantial probability of staying spontaneously in population B is consistent with advanced eusociality in A arising because the evolution of helping efficiency rendered staying spontaneously favored (dashed gray line in Fig. 6A). However, these conclusions are very difficult to draw in practice, particularly because of requirement 2) according to which the population must be naive to the manipulation mechanism evaluated.

## Discussion

Manipulation allows unlikely behaviors to evolve (Dawkins, 1982, Hughes *et al.*, 2012). A puzzle with manipulation is that the evolution of resistance to manipulation can reduce or eliminate the manipulated behaviors (e.g., Parker and Macnair, 1979, Clutton-Brock and Parker, 1995, Gavrilets and Hayashi, 2006, Reuter and Keller, 2001, Kawatsu, 2013). However, the benefits and costs of the manipulated behavior can evolve if they have a genetic basis (Charlesworth, 1978, Worden and Levin, 2007, Akçay and Roughgarden, 2011). Benefits and costs of a manipulated behavior can have a genetic basis since they depend on the extent with which the manipulated behavior is expressed. Yet, how the evolution of payoffs can affect the nature and outcome of the conflict is not known. I have shown that the manipulation conflict can disappear as a result of the evolution of payoffs released by manipulation. The reason is that manipulation can simultaneously favor resistance and the efficiency with which the manipulated behavior is expressed. Since the conflict disappears, I refer to the resulting behavior as being induced rather than as being manipulated. The resolution of conflict has implications for our understanding of the evolution of advanced eusociality in particular, and for the evolution of manipulated behavior in general.

### Ancestrally imperfect resistance and costly inefficiency allow for conflict resolution

The conflict can be eliminated if two key factors occur. First, inefficiency at expressing the manipulated behavior must be costly (*c*_ineff_ > 0). When manipulated, an individual has two options for rebelling: it can either refrain from performing the manipulated behavior (i.e., referred to as resistance), or it can perform the behavior inefficiently. The evolution of these two forms of rebellion can be decoupled because one can be costlier than the other. For simplicity, I have assumed that resistance is costless, and have focused on the effect of costly inefficiency. Inefficiency is costly if the cost of being delayed in the maternal patch is larger than the benefit of exploiting the maternal patch (*c*_*d*_ > *b*_*e*_). That is, inefficiency is costly if the fraction of reproductive opportunities missed by having delayed dispersal is greater than the increased probability to reproduce due to exploiting the maternal patch. If inefficiency is costly, then helping efficiency can increase even if resistance is also favored (compare conditions (4b) and (4c)). Acquiescence becomes favored if helping efficiency becomes large enough. Once acquiescence is favored, the conflict disappears. After the conflict is resolved, both inducing and induced individuals favor the induced behavior, even if the cost of inefficiency disappears.

Second, for the conflict to be eliminated, resistance must be initially imperfect. I have assumed that the manipulated behavior is performed entirely by the subjects of manipulation. So, if they resist with full probability, no manipulated behavior is expressed regardless of how hard manipulators try. In consequence, acquiescence can only be obtained if the probability of resistance is ancestrally imperfect (González-Forero and Gavrilets, 2013). Ancestrally imperfect resistance allows induced behavior to be obtained under more lax conditions than spontaneous behavior. The reason stems from the observation that the evolution of helping efficiency can render both acquiescence and spontaneous behavior favored if they are already present ancestrally. Spontaneous behavior is unlikely to be present ancestrally because it is selected against before ecological conditions make it favorable. In contrast, acquiescence is more likely to be present ancestrally because of the absence of selection for resistance before manipulation arises.

Acquiescence is likely to be present ancestrally depending on how manipulation is executed. Before manipulation starts evolving, there is no initial selection pressure for resistance. Hence, if manipulation is ancestrally executed in a way that subjects of manipulation have not evolved the means to detect, ancestrally imperfect resistance can be expected. Subtle forms of manipulation can then be particularly likely to yield induced behavior.

### Major transitions via conflict resolution

The outcome of conflict resolution is consistent with requirements for a major evolutionary transition in general and for advanced eusociality in particular. Major evolutionary transitions involve the evolution of high levels of cooperation and low levels of conflict (Queller and Strassmann, 2009). Conflict resolution yields here an efficient helping behavior that is induced by the mother and over which there is no conflict between inducing and induced individuals. The high helping efficiency corresponds to high levels of cooperation, while the elimination of conflict produces the required low levels of conflict, thereby fulfilling these requirements for a major transition. On the other hand, advanced eusociality involves 1) maternally induced workers, 2) high levels of specialization of workers and reproductives, and 3) relatively minor conflict in workers regarding their helping role (Wilson, 1971, Michener, 1974, Sherman *et al.*, 1991, Crespi and Yanega, 1995, Hölldobler and Wilson, 2009, Bignell *et al.*, 2011). Maternal manipulation results in 1) maternally induced helping, 2) high helping efficiency, and 3) elimination of conflict between inducing and induced individuals, which directly relate to each of the three mentioned characteristics of advanced eusociality. However, the high maternal fertility observed in the specialization of reproductives is not a consequence of the present model (Figs. S5 and S6 in the SI).

Conflict resolution reinterprets the role of parental manipulation in advanced eusociality. The hypothesis of eusociality via parental manipulation indicates that offspring evolve helping behaviors because of parental influence (Alexander, 1974). Parental manipulation is thought to be relevant for primitive eusociality where the small colony sizes allow the mother to coerce offspring into helping (West, 1967, Michener and Brothers, 1974, Brand and Chapuisat, 2012). However, whether or not parental manipulation is relevant for the evolution of advanced eusociality is less clear, because the large colony sizes would make it impracticable for the mother to coerce offspring into helping (Keller and Nonacs, 1993).

There are at least two kinds of predictions available to assess whether or not parental manipulation occurs in advanced eusocial taxa. A first kind of prediction assumes that the manipulation conflict results in arms races. Predictions of the first kind indicate that the manipulation mechanism (e.g., queen pheromones, and cuticular hydrocarbons) should evolve fast, be highly divergent among species (Brunner *et al.*, 2011), and should not honestly signal the queen’s condition (Keller and Nonacs, 1993). A second kind of prediction assumes that there is a single winner of the manipulation conflict. Predictions of the second kind indicate that if the mother wins the conflict, the maternal preference is satisfied and the fraction of rebellious workers (e.g., those activating their ovaries) should be independent of sister-sister relatedness because the mother is equally related to her female offspring. In contrast, if offspring win the conflict, the offspring preference is satisfied and the fraction of rebellious workers should covary with sister-sister relatedness (Wenseleers *et al.*, 2004, van Zweden *et al.*, 2013). The empirical evidence has not been conclusive, but parental manipulation is only weakly supported in some species (Heinze and d’Ettorre, 2009, Brunner *et al.*, 2011, van Zweden *et al.*, 2013).

The assumptions of the available predictions for testing whether or not parental manipulation occurs in advanced eusociality do not apply if the manipulation conflict is eliminated. After the conflict disappears, evolutionary arms races between inducing and induced individuals are not expected. Instead, the mother and offspring agree on the offspring’s helping role, and should thereafter coevolve in a mutualistic manner. In addition, after the conflict disappears, there is not a single winner of the conflict in the sense of whose preferred outcome is more satisfied, because in this sense both parties win. The resolution of conflict aligns the fitness interests of mother and offspring and both attain their maximum inclusive fitness for their current circumstances. The fraction of rebelling workers after the manipulation conflict is resolved may thus covary with sister-sister relatedness since workers are still able to pursue their own inclusive fitness interests. In addition, large colony sizes are compatible with ancestral manipulation because after the conflict is resolved the mother need not coerce offspring into helping. However, conflict may arise again if the mother evolves multiple mating as it may increase her productivity (Mattila and Seeley, 2007). Finally, the prediction that manipulation mechanisms should not constitute honest signals is not expected after conflict resolution since it is possible that ancestral manipulation is co-opted into honest signaling after the conflict is eliminated (see below).

### Manipulation could either be lost or be co-opted as communication after conflict resolution

Conflict resolution could either eliminate selection for manipulation or it could co-opt manipulation into communication. After conflict resolution, manipulation may become disfavored. Since induced individuals are now favored to express the induced behavior, they may be selected to express it even if manipulation is not present. Suppose that first-brood individuals receive environmental cues (e.g., temperature or humidity) that inform them that they belong to the first brood rather than to the second one. In that case, manipulation becomes unnecessary and first-brood offspring develop as workers following environmental cues. Hence, manipulation could decrease and disappear. Even in the absence of manipulation, the behavior is maintained since the attained helping efficiency renders the behavior favored by selection. However, the behavior is not socially induced anymore, and it becomes environmentally induced instead.

Alternatively, manipulation may continue to be favored after the conflict is eliminated. Now suppose that first brood individuals receive no reliable cues to inform them of the brood they belong to. If manipulation is reduced, helpers may develop in the second brood. Since second-brood helpers do not have recipients of their help, the possibility of second-brood helpers makes manipulation still favored in order to prevent second brood helpers. Manipulation is then maintained after the conflict disappears. In this case, manipulation is maintained to inform first-brood offspring about the brood they belong to. Manipulation is thus co-opted as communication.

The co-option of manipulation as communication also suggests a hypothesis for the evolution of royal jelly in honey bees. In honey bees, royal jelly is given to individuals which induces them to develop into reproductives. That is, individuals are induced to become reproductive rather than workers. The existence of royal jelly is puzzling because individuals should attempt to become reproductive by default. Indeed, it is further puzzling that royal jelly enhances the reproductive abilities of *Drosophila* females (Kamakura, 2011). Why are not these enhanced reproductive abilities in *Drosophila* females present in nature? If manipulation informs offspring about the brood they are in, it may become cheaper to inform reproductives rather than workers. In particular, if the mother starts to produce more workers than reproductives, it may become less expensive to inform reproductives-to-be rather than workers-to-be because there are fewer reproductives. In such a case, induction of reproductives rather than workers would be selected. A mechanism such as royal jelly could then evolve. If reproductives become highly specialized so that they require helpers to survive, their enhanced reproductive abilities triggered by royal jelly are only of use if helpers are available. Then, the enhanced reproductive abilities in *Drosophila* females would be useless in the solitary species.

### Assessing whether a behavior stems from ancestral manipulation

Two analytical conditions specify when a behavior can result from the resolution of manipulation conflict but not from spontaneous behavior. First, the ancestral resistance probability must be sufficiently small (condition (6a) is met). Second, spontaneous behavior must be ancestrally disfavored or, more generally, its ancestral probability must be sufficiently small (condition (S15b) in the SI is not met). Although ancestral conditions cannot be directly estimated except in experimental evolution, indirect estimation of ancestral conditions in extant populations may be possible (Fig. 6B). In addition, estimation of costs, benefits, and relatednesses is very difficult in practice. However, the model defines costs and benefits in a specific manner which may help address this difficulty.

The model presented here is deliberately simple so that complete analytical treatment is possible. Enhancing its realism necessarily affects many of its properties. For example, I assumed that manipulation and resistance are costless. However, costs of manipulation and resistance that are either constant or functions of manipulation and resistance can qualitatively change the dynamics (Reuter *et al.*, 2004). When comparing induced and spontaneous behavior, I assumed that the ancestral benefit *b*_0_ and the ancestral cost *c*_0_ are the same under both scenarios. Yet, the ancestral helping efficiency can be different between these scenarios because individuals may help more or less depending on whether or not and how they were manipulated. I also assumed competition to be global, so the effects of local competition in the conflict resolution remain to be elucidated. In addition, I ignored the effect of genetic drift, which can take the evolutionary trajectories out of the basin of attraction toward induced behavior. Finally, I assumed that the mother manipulates both sexes equally and that both sexes are equally efficient. Although sexually unbiased manipulation and sexually unbiased efficiency are realistic assumptions for diploid genetic systems with ancestral biparental care, they are not proper assumptions for haplodiploids where only maternal care is expected to occur ancestrally. An extension of the model to include sex-differential manipulation and sex-differential efficiency is more appropriate to assess conflict resolution in haplodiploids.

### Conflict resolution in broader contexts

Conflict resolution may similarly occur in other settings where manipulation and resistance coevolve. The model was built for a specific mother-offspring setting so that dynamic analysis is possible. However, the key factors of the process are independent of the mother-offspring setting. Manipulation, resistance, and the efficiency of the manipulated behavior are properties that occur across biological and cultural systems. The necessary factors for conflict elimination, namely ancestrally imperfect resistance and inefficiency costs, can occur widely in evolving systems as well.

Although the manipulation conflict in this model only resolves if the subjects of manipulation and the targets of the manipulated behavior are related (in the model, the “targets” are second-brood offspring; see condition (7b)), the process is in principle not limited to family settings. The conflict may also resolve if subjects and targets are unrelated for at least three reasons. First, relatedness may be unnecessary if resistance is costlier than acquiescence (González-Forero and Gavrilets, 2013). Second, in the model, relatednesses measure the correlation in the heritable components of the traits between actors and recipients of the traits (Frank, 1998, 2013). These correlations may arise from at least six different processes, only one of which requires a family setting. Those processes are: 1) kinship (as in kin selection) (Hamilton, 1964, 1970); 2) conditional response to partner’s behavior (e.g., help only if helped; as in reciprocity) (Queller, 1985, Frank, 1994, Fletcher and Zwick, 2006); 3) biased assortment among groups (e.g., helpers being more common in some groups than in others; as in group selection) (Queller, 1985, Fletcher and Doebeli, 2009); 4) manipulation (e.g., by changing partner’s behavior to match yours); 5) punishment (e.g., by changing payoffs so that the partner changes its behavior); or 6) partner choice (e.g., by changing partner) (Queller, 2011). Third, if relatedness is negative, induced behaviors that harm the targets of the induced behavior could be obtained (which may be modeled by letting *b*_max_ < 0, causing *b* < 0) (González-Forero and Gavrilets, 2013).

The resolution of conflict as a result of the evolutionary process released by manipulation itself renders manipulation both more likely to be important in nature and more difficult to detect. Increasing the testability of manipulation becomes then a potentially rewarding challenge.

## Acknowledgments

I thank Sergey Gavrilets, Laurent Lehmann, Danielle Mersch, Charles Mullon, Laurent Keller, Brian O’Meara, Jon Wilkins, Jeremy Auerbach, Kelly Rooker, Simon Powers, and three anonymous reviewers for discussion or comments on various versions of the manuscript which substantially improved its final form. I thank S. Gavrilets for suggesting a cleaner way to derive the basin of attraction toward induced behavior, and for access to the Volos computer cluster in which numerical simulations conducive to the results reported here were run. Volos is funded by the NIH grant #GM-56693 to S. Gavrilets. I was funded by a Graduate Research Assistantship from the National Institute for Mathematical and Biological Synthesis (NIMBioS). NIMBioS is sponsored by the National Science Foundation, the U.S. Department of Homeland Security, and the U.S. Department of Agriculture through NSF Award #EF-0832858, with additional support from The University of Tennessee, Knoxville.

## Appendix Dynamic equations

The time is discrete. The number of individuals in class *i* at the current time step is *N*_*i*_(*t*). The number of individuals in class *i* in the next time step is *N*_*i*_(*t* + 1), which is given by the *i*-th entry in the column vector **N**(*t* + 1) = **WN**(*t*). For simplicity, fathers can be disregarded and it is enough to keep track of mothers only. Letting the class order in the vector **N** be mothers, first brood, and second brood, the transition matrix is

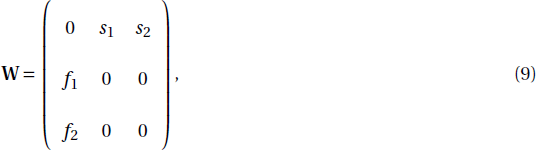

where *s*_*k*_ is the survival of *k*-th-brood offspring (i.e., the probability that *k*-th-brood offspring become mothers) and *f*_*k*_ is the maternal fertility through *k*-th-brood offspring (i.e., the number of offspring produced as brood *k*).

For simplicity, I assume that the fraction of female offspring produced is the same in the first and second broods. Let *σ* be the fraction of offspring that are female. Because for first- and second-brood offspring to become mothers they must be female, then the survival of first-brood offspring is

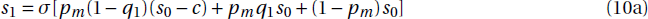

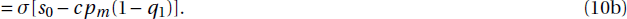

Let *Q* be the average resistance probability among manipulated first-brood offspring in the maternal patch. Then, the survival of second-brood offspring is

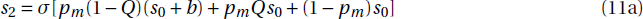

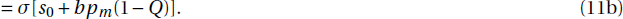

Let *α* be the fraction of offspring that belong to the first brood, and let *n* be the total number of offspring that a mother produces. Each offspring must be weighted by the genetic contribution towards it (Taylor, 1990). The genetic contribution of the mother toward offspring of sex *i* is *η*_*i*_ (i.e., for sexual diploids, *η*_*i*_ = 1/2; for haplodiploids, *η*_♀_ = 1/2 while *η*_♂_ = 1). The genetic contribution of a mother to her offspring is thus on average *η* = *ση*_♀_ + (1 − *σ*)*η*_♂_. Hence, maternal fertility through first and second broods is

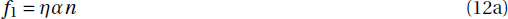

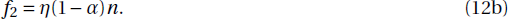

From eq. (29) in Taylor and Frank (1996) and eqs. (6) and (2) in Frank (1997), assuming weak selection and weak mutation, the evolutionary change in the population-average value trait value *z* (= *p*, *q*, *y*) can be approximated by

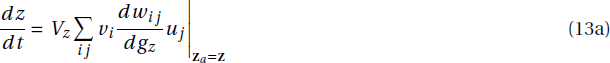

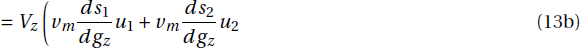

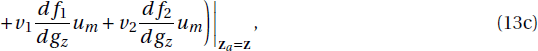

where *w*_*ij*_ is the *ij*-th entry in the transition matrix **W**, *g*_*z*_ is the breeding value for trait *z* in the actor, *V*_*z*_ is the additive genetic variance for trait *z*, *v*_*i*_ is the individual reproductive value for class-*i* individuals, *u*_*j*_ is the equilibrium frequency of class *j* individuals, and traits are evaluated at the population-average value [i.e., **z**_*a*_ = (*p*_*m*_, *q*_1_, *Q*, *y*, *Y*) = **z** = (*p*, *q*, *q*, *y*, *y*)].

Equilibrium class frequencies *u*_*i*_ and individual reproductive values *v*_*i*_ can be respectively obtained from the equations

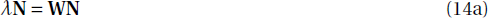

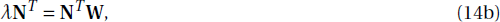

where *T* denotes transposition and the equations are evaluated at the population averages. The equilibrium frequencies *u*_*i*_ are obtained by solving for *N*_*i*_ in eq. (14a) and dividing the solution by ∑*N*_*i*_. The individual reproductive values *v*_*i*_ are obtained by solving for *N*_*i*_ in eq. (14b) together with the condition that the sum of class reproductive values is 1 (i.e., ∑*u*_*i*_*v*_*i*_ = 1, where *u*_*i*_*v*_*i*_ is the reproductive value of class *i*). The quantity *λ* is the dominant eigenvalue of the transition matrix **W**, which gives the asymptotic growth rate of the population. These calculations yield the equilibrium class frequencies

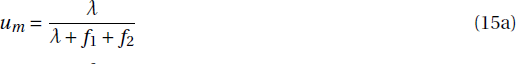

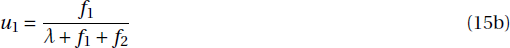

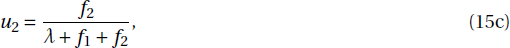

the individual reproductive values

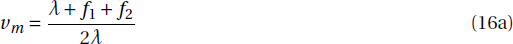

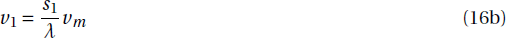

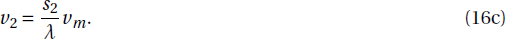

and the asymptotic growth rate

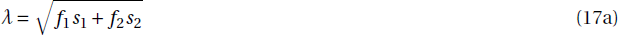

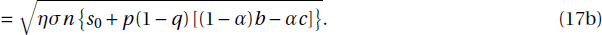

Because the available resources for offspring production only allow the mother to produce a number of offspring that maintains the population size constant, the number of offspring is

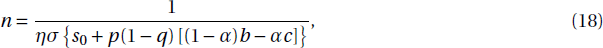

in which case the asymptotic growth rate is *λ* = 1. Since competition is global, the number of offspring *n* depends on the population-average trait values *p*, *q*, and *y* rather than on local average trait values. Hence, because the breeding values of actors are uncorrelated with population averages, the derivatives of fertility in line (13c) are zero.

Therefore, the dynamic equations specified by eq. (13) are

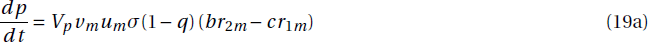

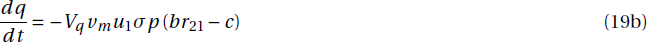

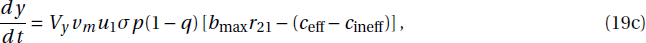

where *r*_*ji*_ = *ρ*_*ji*_*u*_*j*_/*u*_*i*_. The quantity 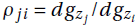 is the regression relatedness of an actor in class *i* toward a recipient in class *j*, where 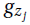 is the breeding value for *z* in the recipient and 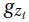 is that in the actor. Hence, *r*_*ji*_ is an equilibrium relatedness.

